# Context-dependent vocal behavior in a glass frog

**DOI:** 10.64898/2026.06.11.731728

**Authors:** Natalia Mejía-Cepeda, Johana Goyes Vallejos

## Abstract

Acoustic communication is fundamental to social interactions in many animal species, allowing individuals to transmit information about identity, reproductive status, and competitive ability. Because call production incurs inherent costs, individuals are expected to modify their vocalizations depending on the social context. However, while context-dependent call variation has been documented in several taxa, including anurans (frogs and toads), glass frogs (Centrolenidae) remain among those for which the acoustic repertoire across social contexts is poorly characterized. Here, we investigated context-dependent call modification in males of the Emerald glass frog, *Espadarana prosoblepon*, comparing calls produced across four social contexts: advertisement in isolation, advertisement in a group, courtship interactions with females, and agonistic interactions with other males. By integrating detailed behavioral field observations with a robust analytical framework, we present evidence that males modify multiple acoustic properties in response to the social context. Specifically, males produced longer, louder advertisement calls when calling in a group than when calling in isolation. Courtship calls contained more notes and were louder than other call types, whereas aggressive interactions were characterized by pulseless, low-frequency, soft calls. Our findings demonstrate that the distinct call types of *E. prosoblepon* are consistently associated with specific social contexts and can be reliably distinguished based on their acoustic structure, providing a framework for future studies investigating the functional significance of context-dependent acoustic signals in anurans.

## Introduction

Acoustic communication is one of the most widespread forms of signaling across the animal kingdom, enabling individuals to convey information about their identity, reproductive status, and competitive ability (Bradbury and Vehrencamp 2011). Acoustic signals, however, are not static; their structure and properties can be actively modified in response to intrinsic and extrinsic factors, including the physical environment, the signaler’s physiological state, and the social context in which communication occurs, such as interactions with rivals or potential mates (Searcy and Nowicki 2010). Among these factors, the social context has been of particular interest in studies of animal communication over the past few decades, because it implies that signalers actively integrate information about their audience to adjust their vocal output, rather than responding to stimuli in a purely hardwired manner (Zuberbühler 2008; Coppinger et al. 2017). Because the social environment in which signals are produced is inherently variable, the ability to modify call structure according to a given social context can be of considerable adaptive value. For instance, signalers are expected to call differently when directing signals toward potential mates versus rival conspecifics (Coppinger et al. 2017). As a result, callers who adjust their signals depending on the social context may improve communication efficiency while mitigating the inherent costs of calling, including energetic investment (Grafe and Thein 2001), increased predation risk (Magrath et al. 2010), and exploitation by unintended receivers (Bernal et al. 2006).

Evidence for socially driven call variation spans a wide range of taxonomic groups, and various studies reveal how specific contexts shape signal production. Audience composition alone can influence signaling. For example, meerkats (*Suricata suricatta*) produce acoustically distinct alarm calls varying in structure according to predator type and urgency level (Manser 2001). Moreover, individuals modify both the type of alarm calls produced and their calling rate depending on whether they are alone versus when in groups, suggesting that alarm signaling is shaped not only by external threats but also by who is listening (Townsend et al. 2012). The identity of that audience can also influence signal plasticity. Male Bengalese finches (*Lonchura striata domestica*) produce songs directed at females that differ in rate, amplitude, and syllable stereotypy compared to those sung in isolation (Sakata et al. 2008), and male brown ghost fish (*Apteronotus leptorhynchus*) produce longer electric signals during courtship than during same-sex interactions (Zakon et al. 2002). Similarly, direct male-male competition also influences acoustic signals. For instance, in katydids of the *Mecopoda* complex, males increase amplitude-modulated signals when females are present but reduce signaling duration during male-male acoustic interactions (Krobath et al. 2017).

In anurans (frogs and toads), males produce advertisement calls—vocalizations that serve the dual function of attracting females and deterring rival males (Wells 1977). However, many species produce additional call types, including courtship and aggressive calls, depending on the social context. Context-dependent acoustic variation has been documented across several well-studied families, including hylids, dendrobatids, and ranids, where males adjust both spectral and temporal call properties in response to rivals, potential mates, or chorus size (Ryan 1988; Welch et al. 1998; Toledo and Haddad 2005; Bee et al. 2010; Fang et al. 2014; Bhat et al. 2022). A recurring pattern is the production of aggressive calls during agonistic encounters between males that differ from advertisement calls in call rate, note duration, number of pulses, and dominant frequency. In the green frog (*Rana clamitans*), for instance, males shift to aggressive calls with a higher rate, longer notes, and lower pitch when responding to a rival (Bee and Perrill 1996); male spring peepers (*Pseudacris crucifer*) similarly switch from tonal advertisement calls to pulsed aggressive calls in the presence of a competitor (Hanna et al. 2014). In some species, these shifts can involve simultaneous spectral and temporal adjustments. For example, in the golden rocket frog (*Anomaloglossus beebei*), males lower the dominant frequency of their calls and increase the number of pulses during aggressive interactions (Pettitt et al. 2012). Similarly, the South American treefrog (*Scinax fuscomarginatus*) produces aggressive calls at lower frequencies and amplitudes than other call types (Toledo and Haddad 2005), and in the neotropical hourglass treefrog (*Dendropsophus ebraccatus*), the introductory note duration increases progressively as competing males draw closer (Reichert 2010). Despite these contributions, context-dependent call variation remains largely unexplored in many anuran groups, such as glass frogs (Centrolenidae), a family in which male vocalizations have been described almost exclusively in the advertisement context (Mendoza-Henao et al. 2021) and for which calls produced during aggressive or courtship interactions remain formally undescribed for the vast majority of species (Dautel et al. 2011; Hutter et al. 2013; Rojas Montoya et al. 2024).

Glass frogs are a group of small, arboreal frogs distributed throughout Central and northern South America, inhabiting tropical and montane rainforests where males typically call from the upper surfaces of leaves overhanging streams (Guayasamin et al. 2009). Currently, more than 170 species are recognized within the family, and male advertisement calls have been described for approximately 60% of them (Mendoza-Henao et al. 2021). However, systematic information on calls produced in other behavioral contexts remains scarce, with fewer than 10% of species having descriptions of vocalizations outside the advertisement context (Dautel et al. 2011; Hutter et al. 2013; Vargas-Salinas et al. 2014; Rojas Montoya et al. 2024). Despite this, field observations suggest that glass frogs engage in a wider array of social behaviors. In a few species, males exhibit high site fidelity, returning to the same calling perch over multiple nights (Angeli et al. 2015; Gómez-Murcia et al. 2024), and have been observed producing vocalizations during physical confrontations with rivals (Dautel et al. 2011; Hutter et al. 2013), when calling in proximity to neighboring males, and during courtship interactions with females (Rojas-Montoya et al. 2024). Together, these observations suggest that acoustic signals in this family likely convey distinct information across social contexts, and that the vocal repertoire of glass frogs is considerably more complex than currently documented.

Among glass frogs, the emerald glass frog (*Espadarana prosoblepon*) is one of the most widely distributed and relatively well-studied species within the family (Jacobson 1985; Hedman and Hughey 2015; Gómez-Murcia et al. 2024; Goyes Vallejos et al. 2024; Rodríguez-Correa et al. 2024), making it an excellent system for investigating context-dependent acoustic variation. Males typically call at night from fixed locations in vegetation near streams and creeks, often maintaining the same calling site both within and across nights (Robertson et al. 2008), and displaced individuals have been shown to return consistently to their specific calling sites (Gómez-Murcia et al. 2024). While the advertisement call has been described in detail, reports of males calling during aggressive encounters exist only as brief field reports (Krohn and Voyles 2014; Hedman and Hughey 2015; Rios-Soto et al. 2017), and the acoustic characteristics of calls produced outside the advertisement context have not been formally described.

Our understanding of the natural history of *E. prosoblepon*, combined with field reports of vocalizations outside the advertisement context, led us to hypothesize that this species’ vocal repertoire includes acoustically distinct call types associated with different social contexts. To test this, we focused on a population of *E. prosoblepon* from southwestern Costa Rica, and we examined how males modify the structure and temporal patterns of their calls across four social contexts: advertisement calling in isolation, advertisement calling within a group of conspecific males, aggressive calling during physical encounters with rivals, and courtship calling in the presence of a nearby female. By integrating field-based behavioral classifications with multivariate analyses of acoustic properties and generalized linear mixed models to quantify differences in individual acoustic parameters, we assessed whether these contexts correspond to acoustically distinguishable call types.

## Methods

### Study site

Our study was conducted at the Wilson Botanical Garden of Las Cruces Research Station (8°47’10”N, 82°57’32”W, WGS84 datum), operated by the Organization for Tropical Studies in the province of Puntarenas, Costa Rica. The site is situated at an elevation of approximately 1,200 meters above sea level and receives approximately 4,000 mm of annual rainfall. The botanical garden is dominated by heliconias, gingers, and palms, and it is crisscrossed by various creeks and ditches that provide suitable breeding habitats for *E. prosoblepon.* Fieldwork was conducted in June and July 2024 and 2025, coinciding with the onset of the rainy season (May–November).

### Acoustic recordings across social contexts

We obtained 107 recordings from 89 *E. prosoblepon* males between 1900 and 2230 h in two 50-m transects along “Culvert Creek,” using a Tascam DR-40X recorder (44.1 kHz sample rate, 16-bit resolution) and a handheld Sennheiser MKE 600 directional microphone (sensitivity: 21 mV/Pa, frequency range: 40 Hz – 20 kHz) at a distance of 50 cm from the focal male. Sound pressure level was also measured at 50 cm, using an Extech 407730 sound level meter (precision of ±2 dB). Recordings varied in length, with individuals producing between one and 15 calls per recording (median = 5). Ambient temperature and humidity at the male’s calling position were measured using a Kestrel 3000 pocket weather meter. Each male was marked with a Visible Implant Elastomer (Northwest Marine Technology, Inc.) and photographed for individual identification to prevent re-recording the same individual in the same social context.

Based on the behavioral context in which males were observed vocalizing, recordings were *a priori* classified into four categories: calls produced by males vocalizing without other calling males in their immediate vicinity were classified as “advertisement call (single)” (*N* = 30). Calls recorded when a focal male was vocalizing in the presence of two to ten conspecific males within two meters were classified as “advertisement call (group)” (*N* = 30). Male vocalizations produced within 0.5 meters of a female and preceded amplexus formation (mating embrace) were defined as “courtship” (*N* = 30). Lastly, acoustic signals emitted during physical agonistic encounters between males were categorized as “aggressive” (*N* = 17) (Figure 1a–d). All recorded aggressive interactions involved two males, each already engaged in physical combat when found, often hanging upside down from a leaf, as shown in Figure 1d. During these encounters, only one of the two males produced aggressive calls, and this was the individual we recorded. Courtship calls were the only calls experimentally induced to be recorded, except in three cases in which males were found vocalizing near a female before initiating amplexus. We searched for pairs in amplexus and gently separated the male and female, positioning them approximately 10 cm apart. This procedure reliably elicited courtship calls immediately before males attempted to resume amplexus.

**Figure 1.**
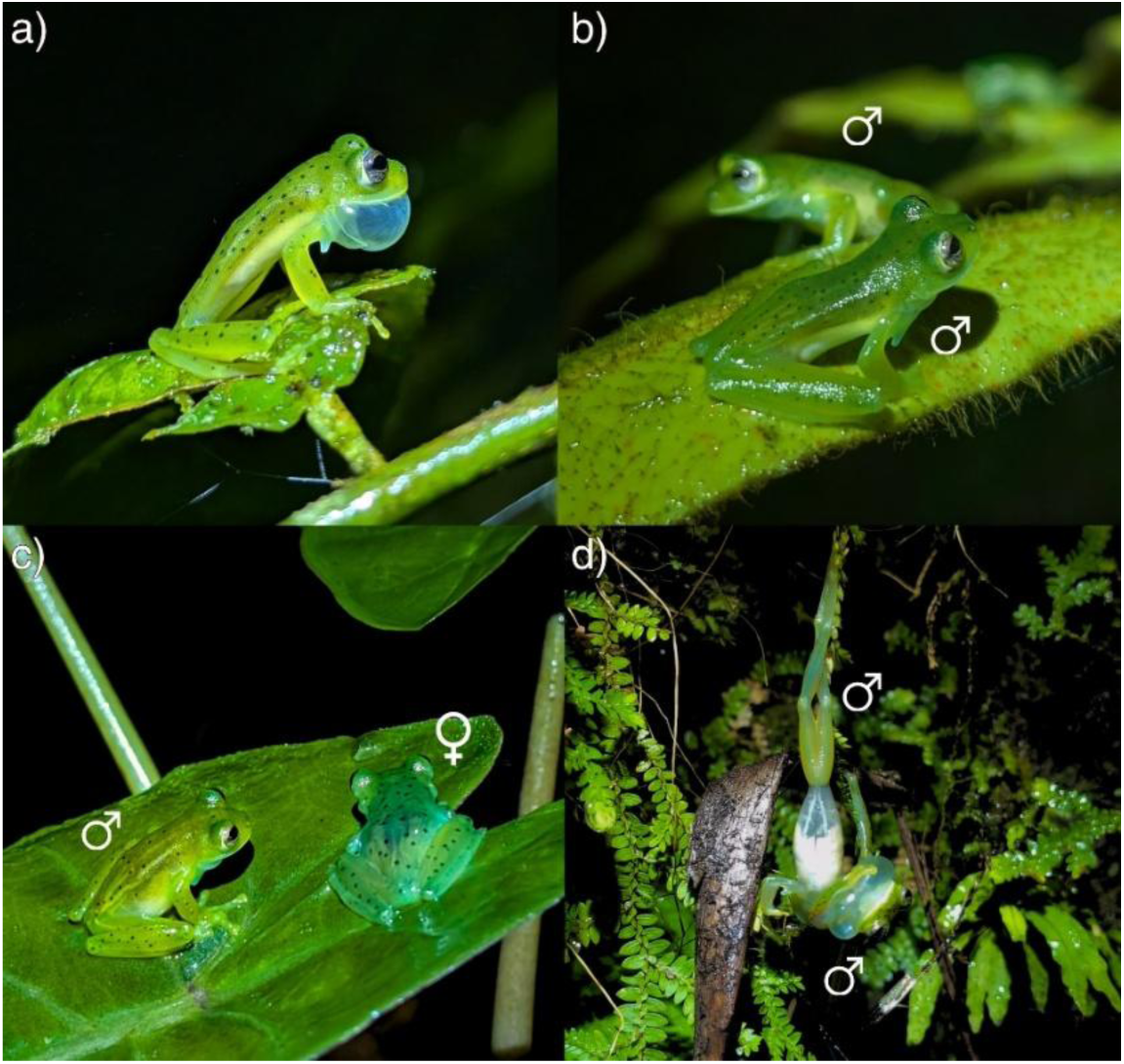
Social contexts used to classify male vocalizations in *E. prosoblepon*: (a) advertisement (single); (b) advertisement (group); (c) courtship; and (d) aggressive.

All the behavioral observations and individual manipulations in the field followed the ASAB/ABS guidelines for the ethical treatment of animals in behavioral research and teaching. Our study was approved by the Costa Rican Ministry of the Environment and Energy (MINAE) and National System of Conservation Areas (SINAC) (approval numbers: R-SINAC-PNI-ACLAP-029-2024 and SINAC-ACLAP-DR-GASP-PNI-R-0019-2025), as well as the Animal Care and Use Committee (ACUC) of the University of Missouri (Protocol No. 45158).

### Acoustic properties dataset

For the acoustic description of each call type, we used a note-centered approach (Köhler et al. 2017; Figure 2) and quantified eight acoustic properties describing the temporal and spectral structure of each call: 1) call duration (s), 2) inter-call interval duration (s), 3) number of notes per call, 4) note duration (s), 5) inter-note interval duration (s), 6) note repetition rate (notes/s), 7) pulses per note, and 8) dominant frequency (Hz). Recordings were visualized and measured in Raven Pro v1.5 (Cornell Laboratory of Ornithology, Ithaca, NY) using a Hanning window function and a window length of 1024 samples. Temporal variables were measured from oscillograms, and the dominant frequency was determined from spectrograms. Because the number of calls per recording varied among males, we summarized the acoustic properties at the individual level. For each male, we first averaged properties measured at the note level (i.e., pulses per note, note duration, and inter-note interval duration) within each call, and then averaged these values across all calls to obtain a single mean value per individual. For properties measured at the call level (call duration, inter-call interval duration, number of notes, and dominant frequency), we averaged values directly across all calls for each male. We also incorporated sound pressure level (dB) as an additional acoustic property in this dataset; this measurement was taken once per recording. Before subsequent statistical analyses, we visually inspected the dataset using box plots to identify extreme values and general patterns within each call type. One clear outlier (an extreme inter-call interval value for individual M85, call type advertisement (single)) was removed before analyses. The ambient temperature during sampling was relatively constant (mean 20.43 ± 0.97 °C; range 18.3 – 22.7 °C; relative humidity = 95%), and thus, temperature effects on call properties were considered negligible.

**Figure 2.**
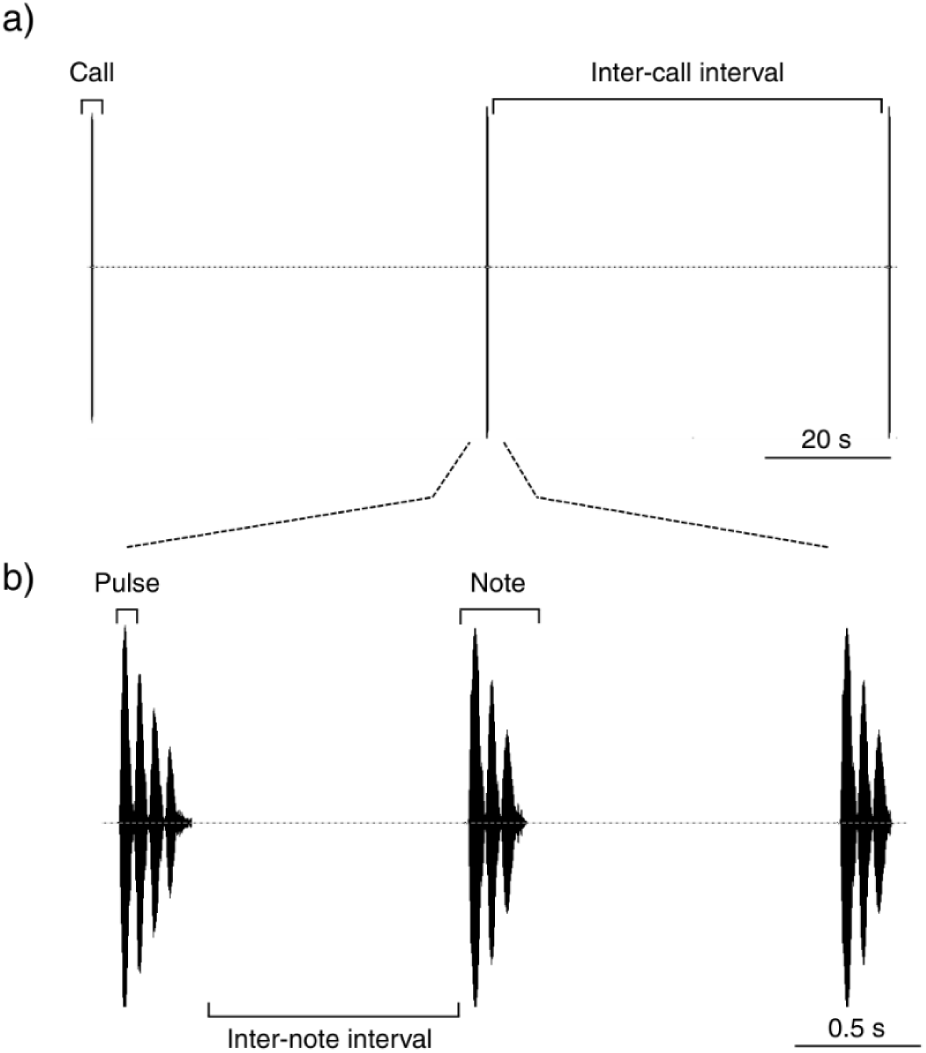
Schematic representation of the note-centered approach used for the quantification of the acoustic properties of the calls of *E. prosoblepon*. (a) Oscillogram showing a sequence of three calls separated by inter-call intervals. (b) Zoomed-in view of a call, illustrating notes, pulses, and inter-note intervals.

### Multivariate Analyses

To evaluate the validity of our *a priori* classification of *E. prosoblepon* call types, we conducted two complementary multivariate analyses—a Cluster Analysis (CA) and a Discriminant Function Analysis (DFA)—performed in R (version 4.5.2; RStudio Team 2025) using RStudio (version 2025.9.2.418; RStudio Team, 2025). To reduce dimensionality and generate a dataset for these analyses, we performed a Principal Component Analysis (PCA) using all nine acoustic properties. Before running the PCA, mean values per individual were standardized to z-scores (mean = 0, SD = 1) to ensure comparability across units (e.g., Hz, dB, seconds). Because the PCA cannot be performed with missing values, we addressed missing data using the *missMDA* package, which imputes missing values via the ‘imputePCA’ function (Josse and Husson 2016). We imputed values for one removed outlier and for missing inter-call interval duration values for individuals with only one recorded call, as this property requires at least two calls to calculate. Imputing these values allowed us to retain this property. We then performed the PCA on the imputed dataset using the ‘prcomp’ function in base R. We used the first four principal components (PC1 – 4) for the subsequent Cluster and Discriminant Function analyses.

The CA was used to identify natural groupings among call types based on the similarity of their acoustic properties, without relying on prior information on call classification. We estimated the optimal number of clusters (*k*) using the *NbClust* package (Charrad et al. 2014), evaluating *k* values from 2 to 10, and applying the *k*-means clustering method with Euclidean distance. The final *k* was selected by majority rule across the 30 validity indices returned. In addition, we performed a DFA, a supervised classification method. The DFA identifies the linear combination of variables that best separates predefined groups. Using cross-validation, the DFA generated posterior classifications based on the acoustic properties, and classification accuracy was quantified as the proportion of males correctly reclassified to their original call type. To account for agreement expected by chance and potential effects of unequal call type sample sizes on classification probabilities, we calculated Cohen’s Kappa coefficient, which quantifies the level of agreement between the observed and predicted classifications adjusted by chance (Titus et al. 1984). Cohen’s Kappa values range from -1 to 1, where values closer to 1 indicate strong agreement, and negative values indicate classification disagreement.

We also present a summary of the acoustic variables measured for each call type in Table 1, excluding males that were not accurately classified by both the CA and DFA, following Pettitt et al. (2012). For continuous acoustic properties, we report the mean ± SD and range; for discrete acoustic properties, we report the median, interquartile range, and range.

**Table 1.**
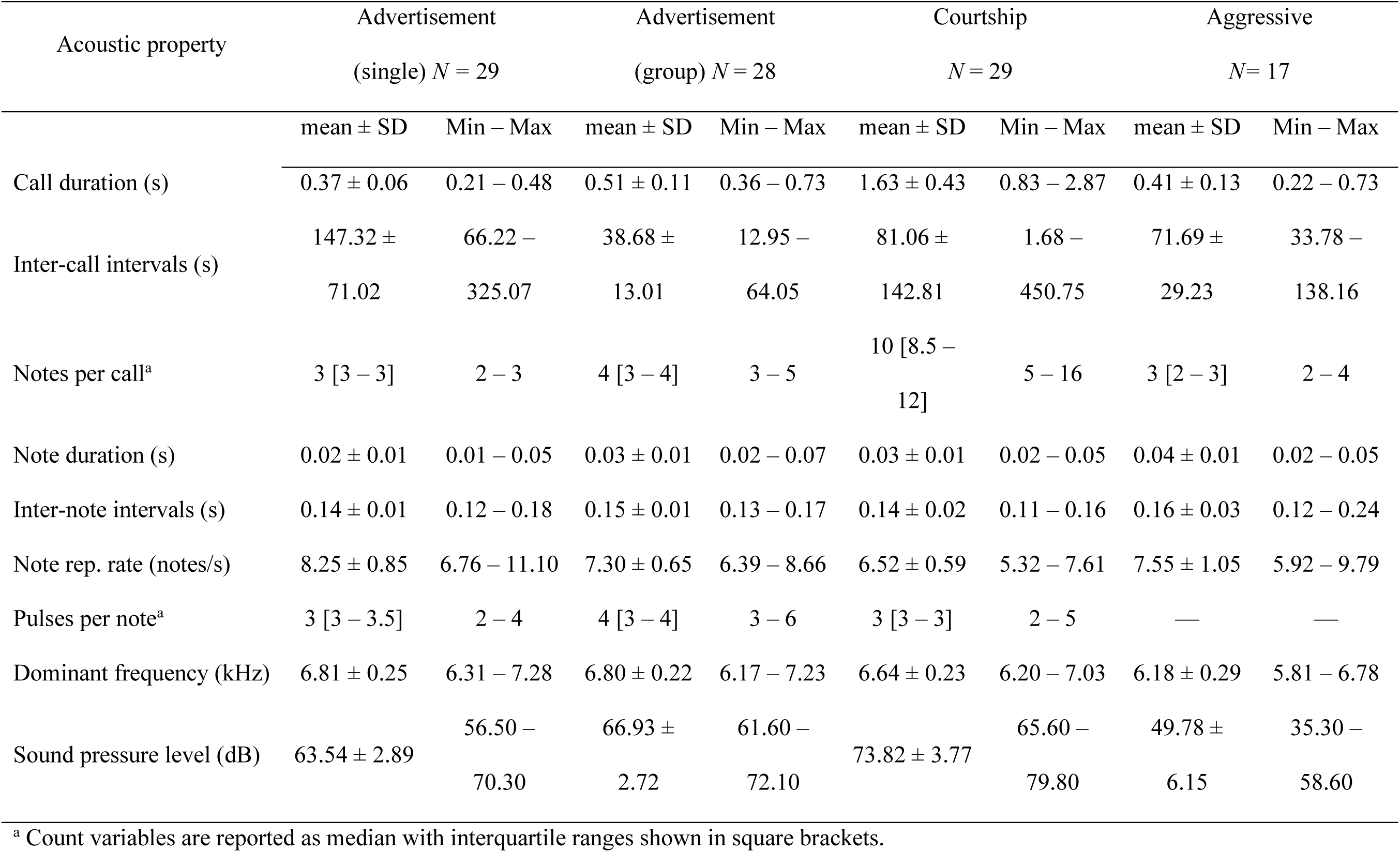
Descriptive statistics for the nine acoustic properties measured from male calls of *E. prosoblepon.* Values summarize the dataset by first averaging measurements within each call type for each male, then averaging these values across individuals.

### Differences in acoustic properties across call types

To assess how the acoustic properties differ among *E. prosoblepon* call types, we conducted separate generalized linear mixed models (GLMM) with each acoustic property as a function of call type (advertisement (single), advertisement (group), courtship, and aggressive call), with ‘Recording’ specified as a random intercept to account for within-individual variation, using the full dataset for all the males accurately classified in the multivariate analyses. All models were fitted using the *glmmTMB* R package (Brooks et al. 2017), and residual diagnostics were checked using the *DHARMa* package (Hartig et al. 2024). For each acoustic property, we selected the error distribution that best fitted its data type. We used a Poisson distribution for the number of notes per call and the number of pulses per note. For the analysis of the number of pulses per note, aggressive calls were excluded because pulsed notes were rare and highly unbalanced across individuals. Only seven males (out of 17) produced any pulsed notes, and among these males, not all aggressive calls were pulsed (median total number of aggressive calls per individual = 5, range = 3–7; median number of pulsed calls = 3, range = 1–5). Call duration, note duration, and note repetition rate were right-skewed continuous variables, and the GLMMs were fitted using a gamma distribution and a log link function. For inter-note interval and inter-call duration (both continuous right-skewed variables), we log-transformed the data and used a Gaussian distribution. The dominant frequency followed approximately a normal distribution, and the GLMM was fitted using a Gaussian distribution. Sound pressure level was the only variable modeled with a Gaussian generalized linear model, as we had only a single value per individual and therefore an individual-level random intercept was unnecessary. To interpret and compare model results, we estimated marginal means using the *emmeans* package (Lenth and Piaskowski 2026) and performed Tukey-adjusted pairwise comparisons among call types.

### Call repetition rate in advertisement calls

Call repetition rate in anurans is a fundamental property of advertisement calls that has been shown to correlate with male attractiveness and reproductive success (e.g., Docherty et al. 2000; Wells and Schwartz 2007). Because courtship and aggressive calls occur in brief, discrete bouts (typically immediately before amplexus or during agonistic encounters), a sustained call rate cannot be meaningfully calculated for those contexts. Therefore, we restricted comparisons of call repetition rate to advertisement calls only (single vs. group). For this, we fitted a GLMM with a gamma distribution and a log-link function, with call repetition rate as a function of call type, with ‘Recording’ as a random intercept. Model diagnostics were checked using the *DHARMa* package (Hartig et al. 2024).

## Results

In this study, we quantitatively analyzed 107 field recordings of male *E. prosoblepon* vocalizations. Males of *E. prosoblepon* produced different acoustic signals depending on the social context in which they gave the call. Based on our field observations and recordings, we categorized these vocalizations into four distinct call types, each consistently associated with a specific behavioral context. When calling alone (single), males produced advertisement calls composed of a sequence of three short notes (Figure 3a); however, in the presence of other conspecific males (group), the advertisement calls typically included an additional note (Figure 3b). Courtship calls were produced when a female approached a calling male. These calls were longer and louder, with males emitting several notes (five or more) before amplexus (Figure 3c). During agonistic encounters, males produced a distinct, soft, and short call consisting of two to four notes, which, in contrast to the other contexts, exhibited lower dominant frequencies and sound pressure levels (Figure 3d).

**Figure 3.**
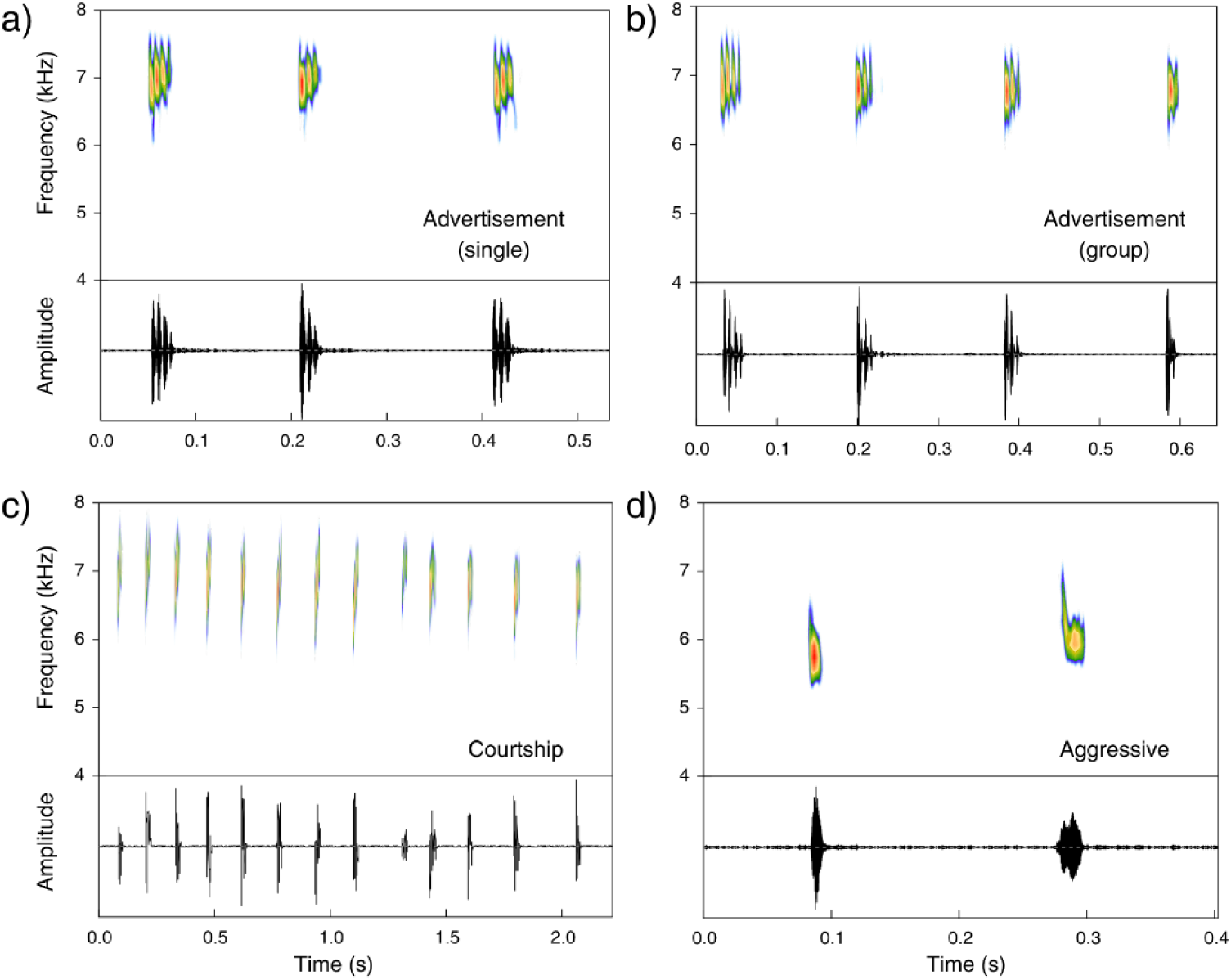
Spectrogram and oscillogram of an (a) advertisement call (single), (b) advertisement call (group), (c) courtship call, and (d) aggressive call of male *E. prosoblepon*.

### Multivariate analyses

For both the CA and DFA, we used the first four components of the PCA, which accounted for approximately 82% of the total variance in the data (Table 2). PC1 accounted for 35.74% of the variation and was positively associated with call duration, number of notes, and sound pressure level. PC2 explained 21.95 % of the variation and was associated with negative loadings for note repetition rate, pulses per note, and dominant frequency. PC3 explained 12.65% of the variation and showed a strong negative loading for note duration, and PC4 explained 11.57% of the variation and had a strong negative loading for inter-note interval duration.

**Table 2.** Loadings of the first four principal components, with values greater than 0.4 highlighted in boldface.

| Acoustic property | PC1 | PC2 | PC3 | PC4 |
| --- | --- | --- | --- | --- |
| Call duration (s) | **0.50** | 0.23 | 0.21 | 0.13 |
| Inter-call intervals (s) | -0.11 | -0.15 | 0.21 | -0.04 |
| Notes per call | **0.50** | 0.19 | 0.26 | 0.21 |
| Note duration (s) | -0.01 | 0.26 | **-0.80** | 0.30 |
| Inter-note intervals (s) | -0.17 | 0.37 | 0.03 | **-0.78** |
| Note repetition rate (notes/s) | -0.37 | **-0.40** | 0.26 | 0.32 |
| Pulses per note | 0.28 | **-0.45** | -0.32 | -0.20 |
| Dominant frequency (kHz) | 0.15 | **-0.53** | -0.17 | -0.26 |
| Sound pressure level (dB) | **0.48** | -0.22 | -0.01 | -0.18 |
| Variance % | 35.74 | 21.95 | 12.65 | 11.57 |
| Cumulative variance % | 35.74 | 57.69 | 70.34 | 81.91 |

The CA validity indices identified three clusters as the optimal number of groupings (*k*) based on similarities in the acoustic properties of male calls (Figure 4). Recordings of advertisement calls (single) and advertisement calls (group) were grouped into a single cluster (cluster 1). Recordings of courtship calls were grouped into cluster 2, while aggressive calls were grouped into cluster 3. The CA misclassified three individuals: two males (M42 and M64), who were originally recorded producing advertisement calls within a group, were assigned to cluster 2 (courtship calls). Additionally, a male that was recorded producing advertisement calls in isolation (M88) was assigned to cluster 3 (aggressive calls). Male M42 and M64 produced advertisement calls with durations nearly twice the average for this call type (mean ± SD = 0.87 ± 0.18 s and 0.84 ± 0.28 s, respectively), a higher than average number of notes per call (median [IQR] = 5.5 [5 – 6] and 5 [4 – 6], respectively), and high sound pressure level values (71.2 and 69.7 dB, respectively). These acoustic properties loaded strongly on PC1 and contributed to the separation of advertisement, courtship, and aggressive calls, where courtship calls are longer and louder than the other call types. Male M88 exhibited a dominant frequency lower than the minimum of the observed range for advertisement calls (mean ± SD = 6.07± 0.3 kHz), a trait that loaded strongly and negatively on PC2 and contributed to the separation of aggressive calls from other call types, possibly accounting for its misclassification.

**Figure 4.**
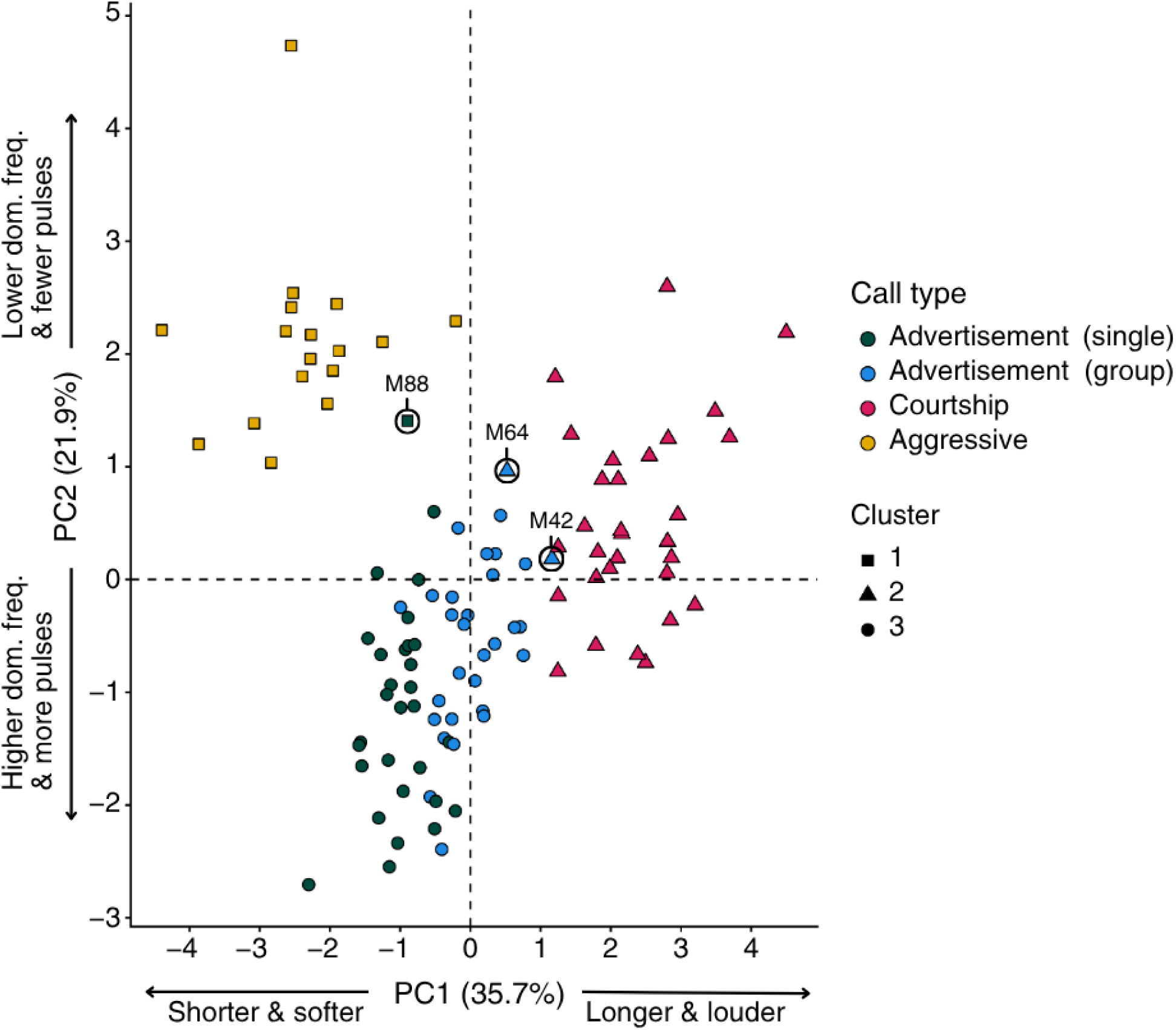
Cluster analysis of PCA scores (PC1 vs. PC2). Points are colored by *a priori* call types and shaped by cluster assignment. PC1 (35.7% of variation explained) describes a gradient from shorter, softer calls to longer, louder calls, whereas PC2 (21.9%) describes a gradient from calls with higher dominant frequencies and more pulses to calls with lower dominant frequencies and fewer pulses. Misclassified males (M88, M64, and M42) are highlighted in black circles.

The DFA correctly classified 97 out of 107 male recordings (91% accuracy; Cohen’s Kappa = 0.87) as belonging to our *a priori* classifications. Classification success varied among call types (Figure 5). Advertisement calls (single) were correctly classified in 25 out of 30 males (83% accuracy), with the remaining five males misclassified as advertisement calls (group). Advertisement calls (group) were correctly assigned for 26 out of 30 males (87 % accuracy), while four males were misclassified as advertisement calls (single). Courtship calls were correctly classified in 29 out of 30 males (97% accuracy), with one male recording (M66) misclassified as an advertisement call (group). This individual produced a shorter courtship call with fewer notes (1.26 s, 7 notes), fewer pulses (median = 2), and a lower sound pressure level (68.3 dB), which may have contributed to the incorrect assignment. All aggressive calls (17 of 17 males) were correctly classified (100% accuracy).

**Figure 5.**
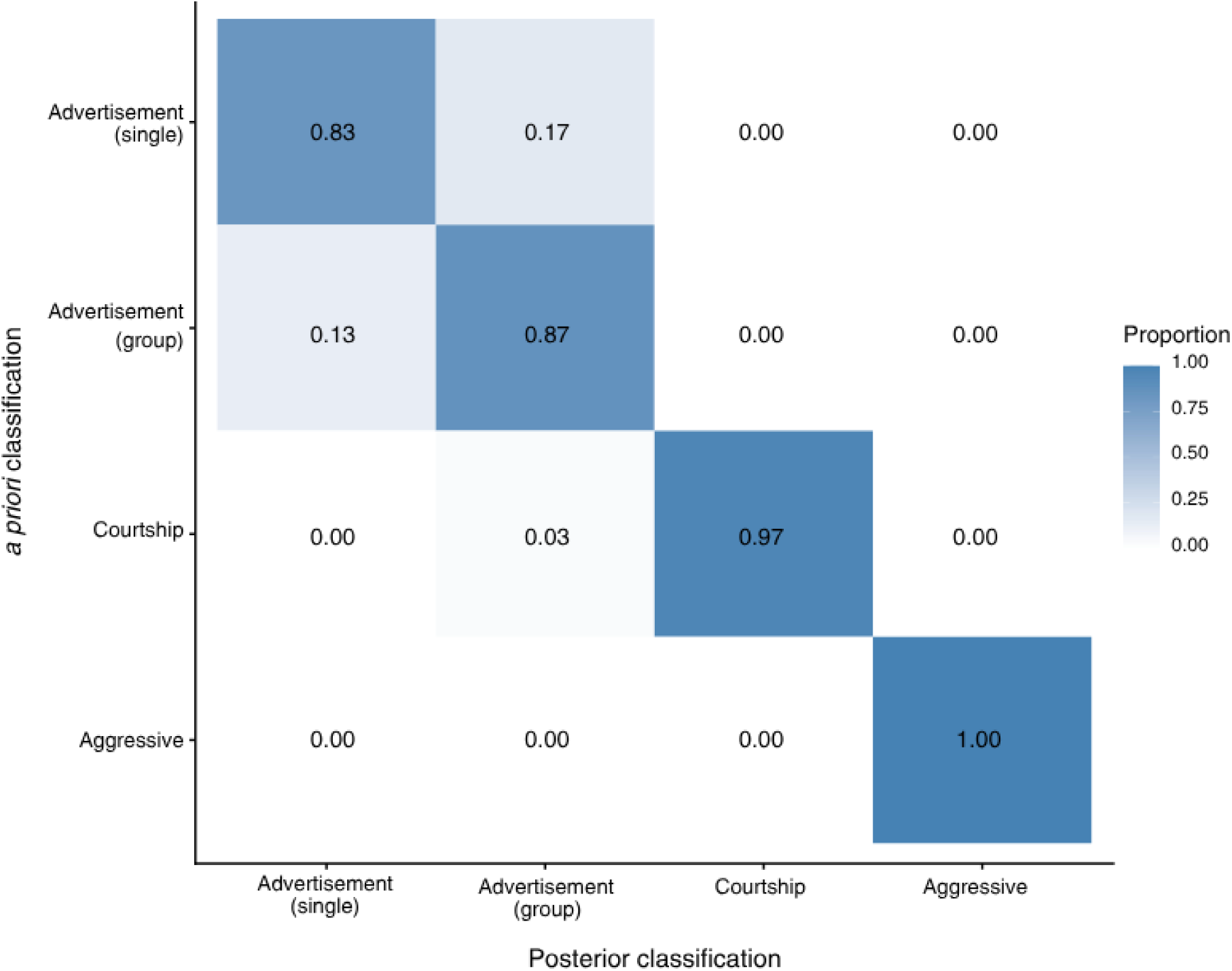
Confusion matrix heat map comparing *a priori* call type classifications with posterior DFA assignments. Cell values represent the proportion of calls classified to each posterior category within each *a priori* call type, with the number of males recorded indicated in parentheses. Darker shading indicates higher classification agreement.

Misclassifications between advertisement calls (single vs. group) were considered biologically plausible because both are produced in the same behavioral context and were therefore retained in their *a priori* classifications. In contrast, misclassifications across other social contexts (i.e., male M66) were excluded from further analyses comparing specific acoustic properties across call types. Based on the combined results of the CA and DFA, males M42, M64, M88, and M66 were removed. The final data set included 29 recordings of male advertisement calls (single), 28 advertisement calls (group), 29 courtship calls, and 17 aggressive calls.

### Differences in acoustic properties across call types

A distinct combination of temporal, spectral, and amplitude features characterized each call type. In terms of call duration, advertisement calls (single) and aggressive calls were similar in length (p = 0.4316); however, this was not true for the other call types. Advertisement calls (group) were significantly longer than advertisement calls (single; p < 0.0001) and aggressive calls (p = 0.0117), while courtship calls were the longest overall, exceeding the duration of both advertisement types (p < 0.0001; Figure 6a). Inter-call intervals also differed significantly among call types. Advertisement calls (single) had the longest inter-call intervals (p < 0.0001), whereas advertisement calls (group) and courtship calls had the shortest inter-call duration and did not differ significantly from each other (p = 0.9770). Aggressive calls showed intermediate inter-call intervals, differing significantly from both advertisement calls (p = 0.0186 and p < 0.0001 single and group, respectively), and courtship calls (p = 0.0417; Figure 6b).

**Figure 6.**
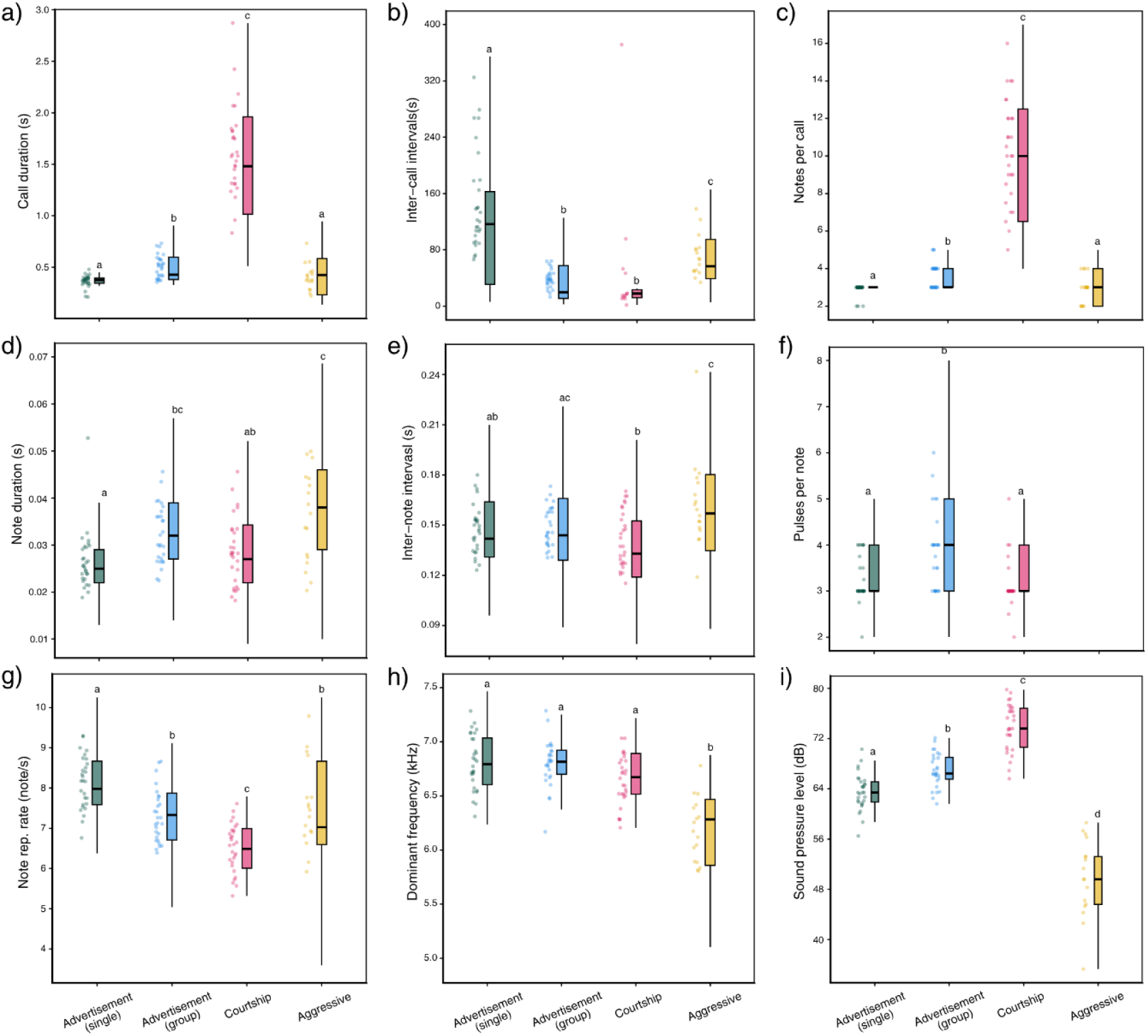
Boxplot showing the nine measured acoustic properties across call types of male *E. prosoblepon*. Boxes indicate the interquartile range with medians shown as horizontal lines, and whiskers extend to the minimum and maximum values. Points represent the mean values of each acoustic variable calculated for individual males. Different letters above each box indicate statistically significant differences between call types based on pairwise comparisons.

Notes, pulses, and note repetition rate further differentiated call types. The number of notes per call was significantly different across all call types (all p < 0.0001), except between advertisement calls (single) and aggressive calls (p = 0.9287). Advertisement calls (single) consisted of an average of three notes and increased to up to five notes during advertisement calls (group). Courtship calls contained the highest number of notes, reaching up to 16 per call, whereas aggressive calls consisted of two to four notes (Figure 6c). Regarding note duration, aggressive calls had the longest overall, differing significantly from advertisement calls (single; p = 0.005) and courtship calls (p = 0.0078), but not from advertisement calls (group; p = 0.5888). Comparing advertisement calls (single vs. group), advertisement calls (group) had significantly longer notes than advertisement calls (single; p = 0.031), while note duration did not differ significantly between either advertisement call type and courtship calls (p = 0.7578 and p = 0.0666 single and group, respectively; Figure 6d).

Aggressive calls had the longest inter-note intervals, which were significantly longer than advertisement calls (single; p = 0.0426) and courtship calls (p < 0.0001), but not advertisement calls (group; p = 0.0654). Advertisement calls (group) had significantly longer inter-note intervals than courtship calls (p = 0.379) but did not differ significantly from advertisement calls (single; p = 0.9955). Advertisement calls (single) and courtship calls did not differ significantly (p = 0.0716; Figure 6e). Pulse numbers did not differ significantly between advertisement calls (single) and courtship calls (p = 0.6741); however, advertisement calls (group) had more pulses per note than both courtship calls (p < 0.0001) and advertisement calls (single; p = 0.0006; Figure 6f). Note repetition rate significantly differed among all call types (all p < 0.01) except between advertisement calls (group) and aggressive calls (p = 0.8878). Advertisement calls (single) had higher note repetition rates than advertisement calls (group), while courtship calls exhibited the lowest repetition rates (Figure 6g).

The dominant frequency was largely conserved across call types, being similar between the two advertisement call types (p = 0.9970) and courtship calls (p = 0.0709 and p = 0.1103 single and group, respectively). Aggressive calls were distinct in exhibiting significantly lower dominant frequencies than all other call types (p < 0.0001; Figure 6h). Lastly, sound pressure levels differed significantly across all call types. Advertisement calls (group) were louder than advertisement calls (single). Courtship calls were the loudest call type, whereas aggressive calls were the softest (all p < 0.0001) (Figure 6i).

Overall, advertisement calls produced by males alone and those produced near other males differed in most measured acoustic properties, except for inter-note interval duration and dominant frequency. When calling in a group, males produced longer calls at faster rates, with more and longer notes, a greater number of pulses per note, and higher sound pressure levels than when calling in isolation. Courtship calls were the most acoustically distinct, being the longest and loudest call type and containing the greatest number of notes. They were produced with short inter-call intervals but exhibited low note repetition rates, resulting in a call structure that contrasted strongly with both advertisement and aggressive calls. Aggressive calls were similar to advertisement calls produced alone in overall duration and number of notes, but were otherwise acoustically distinct, being short, low-amplitude calls with few notes and the lowest dominant frequencies observed, particularly in contrast to courtship calls.

### Call repetition rate in advertisement calls

To examine how calling behavior changes in the presence of conspecific males, we analyzed call repetition rate separately for both advertisement contexts. Advertisement calls (groups) were produced at a significantly higher repetition rate than advertisement calls (single; p < 0.0001), with males calling nearly four times more frequently in the presence of conspecifics (2.06 ± 0.92 calls/min) than in isolation (0.59 ± 0.23 calls/min; Figure 7). Both advertisement call types defined in this study represent the same behavioral context of mate attraction, yet the presence of other males strongly influenced calling rate.

**Figure 7.**
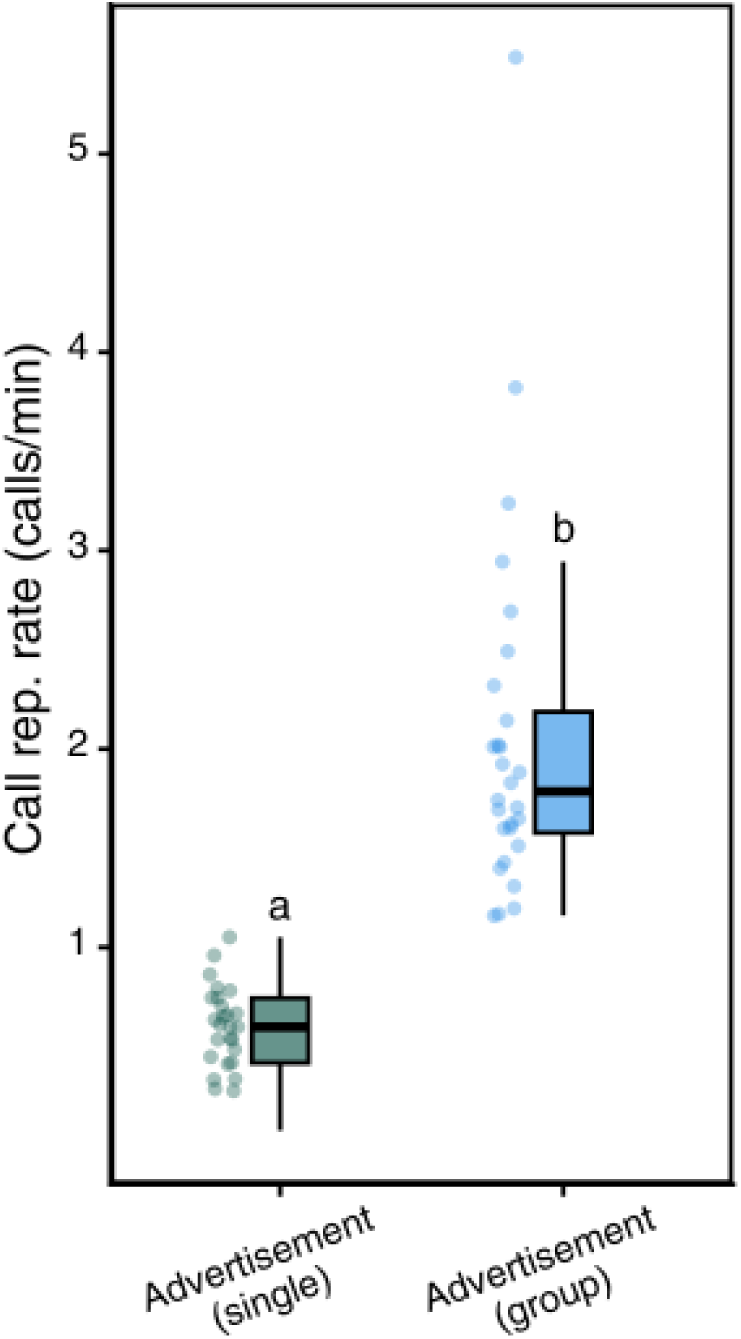
Call repetition rate (calls/min) of advertisement calls (single vs. group) recorded from male *E. prosoblepon*. Boxes indicate the interquartile range with the median shown as a horizontal line; whiskers extend to the minimum and maximum values. Points represent individual male means. Different letters above each box indicate statistically significant differences between call types based on pairwise comparisons.

## Discussion

In this study, we show that males of *E. prosoblepon* exhibit a diverse vocal repertoire, with calls that are consistently associated with distinct behavioral contexts. Building on this, we tested the hypothesis that males modify their acoustic signaling behavior in response to the specific social environment (i.e., calling in isolation, in a group, during aggressive encounters, and in the presence of a female). The multivariate analyses provided a data-driven validation consistent with the behaviors we observed in the field, supporting the conclusion that calls from different behavioral contexts are acoustically distinct. Likewise, generalized linear mixed models examining individual acoustic variables revealed consistent differences in call structure and temporal patterns across behavioral contexts. Advertisement calls are produced in the context of mate attraction and male-male interactions, and the presence of conspecific males clearly influences signal effort: males produce longer, louder, and more calls in a group than when vocalizing alone. Calls produced in a courtship context exhibited the most pronounced structural and temporal divergence, being the longest and loudest, with the greatest number of notes. Conversely, calls produced during physical agonistic encounters exhibited the lowest frequency and sound pressure levels. These findings show that male vocalizations are context-dependent and serve distinct social and reproductive functions.

### Multivariate analyses

Both the CA and the DFA analyses yielded broadly consistent results, grouping the calls into biologically meaningful categories (advertisement, courtship, aggression); however, the DFA outperformed the CA in resolving variation between advertisement calls produced in isolation and in the presence of other males. Cluster analysis is an unsupervised method that groups calls based solely on acoustic similarity, that is, without prior information about behavioral context. Because both advertisement contexts defined separately in this study share a common function (attract mates and deter rivals; Wells 1977) and a similar call structure compared to the other two behavioral contexts, the CA grouped them as a single call type. In contrast, courtship and aggressive calls formed well-defined clusters in the multivariate space, indicating strong acoustic differentiation that, at the same time, reduced separation between the two advertisement contexts. Meanwhile, the DFA is a supervised method that maximizes separation among *a priori* classifications. In our study, the DFA provided higher and finer resolution, effectively distinguishing call categories at a finer scale, particularly for advertisement calls recorded in isolation versus in a group. Over 90% of the calls across our call types were accurately classified, with only a small proportion (<17%) of calls misclassified exclusively to the other advertisement call context (i.e., advertisement calls (single) misclassified as advertisement calls (group) or vice versa). In contrast, the CA misclassified calls across distinct behavioral contexts, such as classifying advertisement calls as courtship or aggressive calls.

In other frog systems, DFA has been shown to be more accurate than CA for classifying acoustic signals. In a similar study that provided a quantitative description of the complete vocal repertoire of the golden rocket frog (*Anomaloglossus beebei*), the DFA slightly outperformed the CA, although both analyses correctly distinguished among three call types (i.e., advertisement, courtship, and aggressive; Pettitt et al. 2012). Remarkably, DFA has also demonstrated strong discriminatory power at the individual level, successfully assigning 91% of calls to the correct individual in the boreal chorus frog (*Pseudacris maculata*; Bee et al. 2010). This pattern has also been reported in other systems that communicate acoustically. In a study comparing the performance of CA and DFA in classifying field cricket (Gryllinae) acoustic signals at the species level, CA achieved moderate clustering success (55–90%), whereas the DFA classified 95–100% of calls correctly, particularly for signal types that exhibited structural overlap (Jaiswara et al. 2013). In echolocating bats, DFA successfully classified 79% of vocalizations at the species level in Britain (Parsons and Jones, 2000) and 80–82% in Italy (Russo and Jones 2002), demonstrating its robustness across different species. In our study, CA accurately grouped *E. prosoblepon* calls into broader, biologically meaningful categories. However, DFA demonstrated greater discriminatory power, enabling us to confirm that there are indeed finer distinctions between advertisement calls produced in isolation versus those produced in a group. In future studies, DFA may even help resolve differences at the individual level. It is important to note that because DFA is a supervised model, its performance relies heavily on accurate *a priori* classifications. Therefore, we underscore the need for detailed natural history observations coupled with acoustic analyses. Without an understanding of the behavioral context in which calls were produced, such fine-scale discrimination would not have been possible.

### Differences in acoustic properties across social contexts

We found that the presence of other males drives differences in all but two acoustic properties (inter-note interval duration and dominant frequency) of *E. prosoblepon* advertisement calls. Under competitive social conditions, males of *E. prosoblepon* increased their overall signaling effort by producing louder, longer calls with additional notes at a higher calling rate than when calling alone. Advertisement calls are among the most energetically costly signals produced by animals (Grafe and Thein 2001), yet increasing call rate, call duration, or call complexity can enhance mate attraction and competitive success (Gerhardt et al. 2000). For instance, in the gray tree frog (*Hyla versicolor*), males calling in dense choruses produced calls that were nearly twice as long and contained more pulses than those of isolated individuals (Wells and Taigen 1986). In the brown-bordered snouted treefrog (*Scinax fuscomarginatus*), males’ calling rate increased proportionally with the number of males in a chorus (Toledo and Haddad 2005). In addition, playback experiments have demonstrated that females of *H. versicolor* prefer long calls when choosing a mate (Gerhardt et al. 2000) or higher calling rates in the case of female marble reed frogs (*Hyperolius marmoratus*; Jennions et al. 1995). Similarly, in two endemic frogs from southwest India (*Nyctibatrachus humayuni* and *Pseudophilautus amboli*), males calling near conspecific competitors produce longer, more complex call sequences than those calling alone, a pattern interpreted as a territorial behavior (Bhat et al. 2022). Thus, although these behavioral changes impose substantial energetic costs, male frogs appear to invest more in advertisement calling when rivals are nearby, as these calls serve not only to attract females but also to mediate male-male interactions and facilitate spacing between calling individuals (Gerhardt 1994; Toledo et al. 2015).

This context-dependent modification of acoustic signals is widespread across animal taxa. In birds, Brumm and Todt (2004) showed that male nightingales (*Luscinia megarhynchos*) increased the amplitude of their songs by more than 5 dB when interacting with a simulated rival, suggesting that producing lower amplitude songs when alone may reduce the energetic costs of singing. Similarly, male saddle-backed bush crickets (*Ephippiger ephippiger*) produce longer and more variable songs within choruses than when calling alone (Ritchie 1992). In mammals, Amichai et al. (2015) demonstrated that bats (*Pipistrellus kuhlii*) extend call duration and increase call amplitude in response to conspecific playbacks, presumably to avoid signal overlap; and male hyraxes (*Procavia capensis*) increase calling rate in response to nearby conspecific male songs (Goll et al. 2017). These examples illustrate a common pattern: individuals strategically adjust their signaling effort in social contexts to optimize communication and navigate competition, despite energetic costs. In our population, males can occur in isolated locations, but along our transects, local density is high, with multiple males calling in proximity and possibly interfering acoustically with one another. Thus, *E. prosoblepon* males appear to strategically adjust their signaling effort in response to these social conditions.

Males optimize call output to enhance communication effectiveness and attractiveness, possibly biasing mate choice toward individuals with higher calling effort while simultaneously mediating spacing and competitive interactions with neighboring males. Two non-mutually exclusive hypotheses may account for the observed variation in call structure: (1) a ‘female attraction hypothesis’, which predicts that longer, louder, or more complex calls increase male attractiveness to females and enhance mating success (Ryan 1988; Gerhardt et al. 2000; Wells and Schwartz 2007), and (2) a ‘male competition hypothesis’, which predicts that these same acoustic modifications function to signal competitive ability to rival males, deterring approach and mediating spacing among calling individuals (Wells 1977; Bradbury and Vehrencamp 2011). Future experiments will be necessary to determine the functional consequences of this context-dependent variation.

Male *E. prosoblepon* courtship calls were the most acoustically distinct vocalizations across the behavioral contexts we described in this study. These calls were typically produced once, immediately preceding amplexus, and were highly stereotyped. In most frog species where male courtship calls have been described, they differ temporally and spectrally from advertisement calls and are generally described as short-range, low-amplitude signals directed specifically at nearby females (Teixeira et al. 2016), often interpreted as subtle signals that may reduce detection by competitors or predators (Toledo et al. 2015). For example, in the smooth guardian frog (*Limnonectes palavanensis*), males produce a soft, single-note courtship call in response to nearby female calls that is significantly shorter than the advertisement call (Goyes Vallejos et al. 2017). Similar short, high-pitched courtship calls have been described in *Pseudophryne covacevichae* and *Austrochaperina robusta* when females approach (Groffen et al. 2024). These signals are thought to either guide females to an oviposition site or signal male receptivity as a female approaches, although their specific function is rarely tested. In contrast to this general pattern, courtship calls in *E. prosoblepon* were the longest and loudest signal in the repertoire, containing approximately three times as many notes as advertisement calls. We categorized this signal as a courtship call because it is produced exclusively in the presence of a female and immediately precedes amplexus. This classification is further supported by the stereotyped sequence of events during mate choice: females approach males at their calling perches, which are not oviposition sites, as egg deposition occurs elsewhere (Goyes Vallejos et al. 2024); females then will typically crouch a few centimeters in front of a male, after which the male produces the courtship call and proceeds directly to amplexus, often continuing to call as he mounts the female. In other vocalizing taxa, courtship calls are known to play important roles in mate assessment and mating decisions. For example, in the field cricket *Teleogryllus oceanicus*, males switch from an advertisement call to a distinct courtship call when females approach. Experimental trials in this system show that females prefer longer courtship calls, which have been interpreted as signals used to assess male quality prior to mating. Satellite males may also exploit these courtship calls to intercept approaching females (Rebar et al. 2009). However, during our field observations, we never observed males of *E. prosoblepon* attempting to intercept or displace an amplectant male, despite their proximity to neighboring males. The function of this courtship call in *E. prosoblepon,* therefore, remains uncertain. It may serve as a final assessment signal directed at the female or as a signal that reinforces female receptivity immediately prior to mating. Alternatively, it could function in deterring nearby males at a critical moment before amplexus. Determining the functional role of this signal will require targeted experimental investigation.

Aggressive calls of male *E. prosoblepon* were recorded during physical agonistic encounters between two males. These calls consisted of low-amplitude signals with a low dominant frequency and were produced during venter-to-venter wrestling while males hung from the vegetation. During our observations, the calls were produced by only one of the males involved in the interaction. Because recordings were made once males were already in physical contact, we cannot determine whether these calls are produced before escalation or occur specifically during combat. Aggressive vocalizations across many animal taxa are often characterized by low-frequency, low-amplitude signals produced at close range during escalated contests, as is the case in *E. prosoblepon*. In frogs, for example, aggressive calls in the snouted tree frog (*Scinax fuscomarginatus*) and the Cerrado gladiator frog (*Bokermannohyla ibitiguara*) have been interpreted as the final stage of a gradual escalation of aggressive behavior culminating in physical combat (Toledo and Haddad 2005; Nali and Prado 2014). Similar patterns occur in other vertebrates. In several families of catfish (Ariidae, Mochokidae, Doradidae, and Pimelodidae), males produce low-frequency sounds during agonistic encounters with conspecifics, and these signals are thought to regulate aggressive interactions by escalating, inhibiting, or modulating conflicts (Ladich 1997). Likewise, male Savannah sparrows (*Passerculus sandwichensis*) produce soft songs during territorial intrusions, and higher singing rates reliably predict attack, suggesting a role in signaling imminent escalation (Moran et al. 2018). However, the social dynamics of aggressive interactions in *E. prosoblepon* remain poorly understood. Previous work has suggested that the male that falls from the leaf during these encounters is considered the loser (Hedman and Hughey 2015), but because we did not observe contests through their conclusion, we cannot determine whether calling males were more likely to be the eventual winners. In fact, the motivation underlying these aggressive interactions also remains unknown. Males may be defending calling sites, as occurs in many anuran species, but the aggressive behavior in *E. prosoblepon* has not been studied in depth. Consequently, the functional significance of the aggressive call remains unexplored. It is also unclear why only one male appears to produce the call during fights, or how this one-sided vocalization influences the dynamics and outcome of contests. Further behavioral and experimental studies will be necessary to clarify the role of aggressive signaling and territoriality in this species.

## Conclusion

Our study demonstrates that the vocal repertoire of the glass frog *E. prosoblepon* is strongly shaped by social context, with males producing acoustically distinct signals during advertisement, courtship, and aggressive interactions. Males modify advertisement calls in the presence of competitors, produce an exaggerated courtship call immediately before amplexus, and emit a distinct aggressive call during male–male interactions. By integrating field behavioral observations with robust acoustic analyses, we show that males flexibly modify multiple acoustic properties, including duration, call rate, dominant frequency, and sound pressure level. These context-dependent patterns suggest that vocalizations in *E. prosoblepon* are shaped by a combination of sexual and social selection, serving specific roles in mate attraction and/or competition. Our classifications are based on observed behavioral contexts rather than experimentally demonstrated signal functions. However, the clear acoustic distinctiveness of each call type allows us to make testable predictions about their functional significance in *E. prosoblepon*. By identifying distinct signals associated with specific social contexts, this study establishes a foundation for future experimental approaches—particularly playback experiments— examining how females and rival males respond to these calls and how acoustic signals mediate mating and competitive interactions in this species.

## Data Availability

Analyses reported in this article can be reproduced using the data provided by Mejía-Cepeda and Goyes Vallejos (2026) https://doi.org/10.5281/zenodo.20560584.

## Acknowledgements

We are grateful to the Organization for Tropical Studies and to the staff at Las Cruces Biological Station in Costa Rica for logistical and scientific support. We also thank Dr. Manuel Leal and Dr. Rex Cocroft for their valuable feedback on the manuscript.

## References

Amichai E, Blumrosen G, Yovel Y. 2015. Calling louder and longer: how bats use biosonar under severe acoustic interference from other bats. Proc R Soc B Biol Sci. 282(1821):20152064. 10.1098/rspb.2015.2064

Angeli NF, DiRenzo G V., Cunha A, Lips KR. 2015. Effects of Density on Spatial Aggregation and Habitat Associations of the Glass Frog *Espadarana (Centrolene) prosoblepon*. J Herpetol. 49(3):388–394. 10.1670/13-110

Bee MA et al. 2010. Assessing Acoustic Signal Variability and the Potential for Sexual Selection and Social Recognition in Boreal Chorus Frogs (*Pseudacris maculata*). Ethology. 116(6):564–576. 10.1111/j.1439-0310.2010.01773.x

Bee MA, Perrill SA. 1996. Responses To Conspecific Advertisement Calls in the Green Frog (*Rana Clamitans*) and Their Role in Male-Male Communication. Behaviour. 133(3–4):283–301. 10.1163/156853996X00152

Bernal XE, Rand AS, Ryan MJ. 2006. Acoustic preferences and localization performance of blood-sucking flies (*Corethrella coquillett*) to túngara frog calls. Behav Ecol. 17(5):709–715. 10.1093/beheco/arl003

Bhat AS, Sane VA, Seshadri KS, Krishnan A. 2022. Behavioural context shapes vocal sequences in two anuran species with different repertoire sizes. Anim Behav. 184:111–129. 10.1016/j.anbehav.2021.12.004

Bradbury JW, Vehrencamp SL. 2011. Principles of Animal Communication. 2nd ed. Sinauer Associates. Brooks ME et al. 2017. Modeling zero-inflated count data with glmmTMB. 10.1101/132753

Brumm H, Todt D. 2004. Male–male vocal interactions and the adjustment of song amplitude in a territorial bird. Anim Behav. 67(2):281–286. 10.1016/j.anbehav.2003.06.006

Charrad M, Ghazzali N, Boiteau V, Niknafs A. 2014. NbClust : An R Package for Determining the Relevant Number of Clusters in a Data Set. J Stat Softw. 61(6):1–36. 10.18637/jss.v061.i06

Coppinger B et al. 2017. Studying audience effects in animals: what we can learn from human language research. Anim Behav. 124:161–165. 10.1016/j.anbehav.2016.12.020

Dautel N et al. 2011. Advertisement and combat calls of the glass frog *Centrolene lynchi* (Anura: Centrolenidae), with notes on combat and reproductive behaviors. Phyllomedusa J Herpetol. 10(1):31–43. 10.11606/issn.2316-9079.v10i1p31-43

Docherty S, Bishop PJ, Passmore NI. 2000. Consistency of calling performance in male *Hyperolius marmoratus marmoratus*: implications for male mating success. African J Herpetol. 49(1):43–52. 10.1080/21564574.2000.9650015

Fang G et al. 2014. Male vocal competition is dynamic and strongly affected by social contexts in music frogs. Anim Cogn. 17(2):483–494. 10.1007/s10071-013-0680-5

Gerhardt HC. 1994. The Evolution of Vocalization in Frogs and Toads. Annu Rev Ecol Syst. 25(1):293–324. 10.1146/annurev.es.25.110194.001453

Gerhardt HC, Tanner SD, Corrigan CM, Walton HC. 2000. Female preference functions based on call duration in the gray tree frog (*Hyla versicolor*). Behav Ecol. 11(6):663–669 10.1093/beheco/11.6.663

Goll Y, Demartsev V, Koren L, Geffen E. 2017. Male hyraxes increase countersinging as strangers become ‘nasty neighbours.’ Anim Behav. 134:9–14. 10.1016/j.anbehav.2017.10.002

Gómez-Murcia DA, Bedoya-Ospina M del M, Arcila-Pérez LF, Vargas-Salinas F. 2024. Nothing like home: most males of *Espadarana prosoblepon* (Anura Centrolenidae) exhibit homing to calling site despite the availability of alternative suitable sites for calling and mating. Ethol Ecol Evol. 36(1):70–85. 10.1080/03949370.2023.2201923

Goyes Vallejos J et al. 2024. Not enough time: short-term female presence after oviposition does not improve egg survival in the emerald glass frog. Anim Behav. 213:161–171. 10.1016/j.anbehav.2024.05.008

Goyes Vallejos J, Ulmar Grafe T, Ahmad Sah HH, Wells KD. 2017. Calling behavior of males and females of a Bornean frog with male parental care and possible sex-role reversal. Behav Ecol Sociobiol. 71(6):95. 10.1007/s00265-017-2323-3

Grafe TU, Thein J. 2001. Energetics of calling and metabolic substrate use during prolonged exercise in the European treefrog *Hyla arborea*. J Comp Physiol B Biochem Syst Environ Physiol. 171(1):69–76. 10.1007/s003600000151

Groffen J, Rush ER, Hoskin CJ. 2024. Calling locations and courtship calls of the frogs *Austrochaperina robusta* Fry, 1912 and *Pseudophryne covacevichae* Ingram and Corben, 1994 in northern Australia. Herpetol Notes. 17:315–321

Guayasamin JM et al. 2009. Phylogenetic systematics of Glassfrogs (Amphibia: Centrolenidae) and their sister taxon *Allophryne ruthveni*. Zootaxa. 2100(1):1–97. 10.11646/zootaxa.2100.1.1

Hanna DEL, Wilson DR, Blouin-Demers G, Mennill DJ. 2014. Spring peepers *Pseudacris crucifer* modify their call structure in response to noise. Curr Zool. 60(4):438–448. 10.1093/czoolo/60.4.438

Hartig F, Lohse L, de Souza Leite M. 2024. DHARMa: Residual Diagnostics for Hierarchical (Multi-Level / Mixed) Regression Models. https://cran.r-project.org/package=DHARMa

Hedman HD, Hughey MC. 2015. Body size, humeral spine size, and aggressive interactions in the emerald glass frog, Espadarana prosoblepon (Anura: Centrolenidae) in Costa Rica. Mesoamerican Herpetol. 2:500–508.

Hutter CR et al. 2013. The territoriality, vocalizations and aggressive interactions of the red-spotted glassfrog, *Nymphargus grandisonae*, Cochran and Goin, 1970 (Anura: Centrolenidae). J Nat Hist. 47(47–48):3011–3032. 10.1080/00222933.2013.792961

Jacobson SK. 1985. Reproductive Behavior and Male Mating Success in Two Species of Glass Frogs (Centrolenidae). Herpetologica. 41(4):396–404. http://www.jstor.org/stable/3892108

Jaiswara R, Nandi D, Balakrishnan R. 2013. Examining the Effectiveness of Discriminant Function Analysis and Cluster Analysis in Species Identification of Male Field Crickets Based on Their Calling Songs Consuegra S, editor. PLoS One. 8(9):e75930. 10.1371/journal.pone.0075930

Jennions MD, Bishop PJ, Backwell PRY, Passmore NI. 1995. Call Rate Variability and Female Choice in the African Frog, *Hyperolius marmoratus*. Behaviour. 132(9/10):709–720. http://www.jstor.org/stable/4535294

Josse J, Husson F. 2016. missMDA : A Package for Handling Missing Values in Multivariate Data Analysis. J Stat Softw. 70(1):1–31. 10.18637/jss.v070.i01

Köhler J et al. 2017. The use of bioacoustics in anuran taxonomy: Theory, terminology, methods and recommendations for best practice. Zootaxa. 4251(1):1–124. 10.11646/zootaxa.4251.1.1

Krobath I, Römer H, Hartbauer M. 2017. Plasticity of signaling and mate choice in a trilling species of the Mecopoda complex (Orthoptera: Tettigoniidae). Behav Ecol Sociobiol. 71(11). 10.1007/s00265-017-2381-6

Krohn AR, Voyles J. 2014. A short note on the use of humeral spines in combat in *Espadarana prosoblepon* (Anura: Centrolenidae). Alytes. 31(3–4):83–85.

Ladich F. 1997. Agonistic behaviour and significance of sounds in vocalizing fish. Mar Freshw Behav Physiol. 29(1–4):87–108. 10.1080/10236249709379002

Lenth R V, Piaskowski J. 2026. emmeans: Estimated Marginal Means, aka Least-Squares Means. https://rvlenth.github.io/emmeans/

Magrath RD, Haff TM, Horn AG, Leonard ML. 2010. Calling in the Face of Danger: Predation Risk and Acoustic Communication by Parent Birds and Their Offspring. 1st ed. Vol 41 Elsevier Inc. 10.1016/S0065-3454(10)41006-2

Manser MB. 2001. The acoustic structure of suricates’ alarm calls varies with predator type and the level of response urgency. Proc R Soc London Ser B Biol Sci. 268(1483):2315–2324. 10.1098/rspb.2001.1773

Mejía-Cepeda N, Goyes Vallejos J. 2026. Data from: Context-dependent vocal behavior in a glass frog. Behav. Ecol. 10.5281/zenodo.20560584

Mendoza-Henao AM, Duarte-Marin S, Rada M. 2021. Advertisement calls of six glassfrog species in the Colombian Andes, and comments on priorities for future research and conservation. Amphib Reptil Conserv. 15(2):156–171. 10.5281/zenodo.16822085

Moran IG et al. 2018. Quiet violence: Savannah Sparrows respond to playback-simulated rivals using low-amplitude songs as aggressive signals. Ethology. 124(10):724–732. 10.1111/eth.12805

Nali RC, Prado CPA. 2014. The fight call of *Bokermannohyla ibitiguara* (Anura: Hylidae): First record for the genus. Salamandra. 50(3):181–184

Pettitt BA, Bourne GR, Bee MA. 2012. Quantitative acoustic analysis of the vocal repertoire of the golden rocket frog (*Anomaloglossus beebei*). J Acoust Soc Am. 131(6):4811–4820. 10.1121/1.4714769

Rebar D, Bailey NW, Zuk M. 2009. Courtship song’s role during female mate choice in the field cricket *Teleogryllus oceanicus*. Behav Ecol. 20(6):1307–1314. 10.1093/beheco/arp143

Reichert MS. 2010. Aggressive thresholds in *Dendropsophus ebraccatus*: habituation and sensitization to different call types. Behav Ecol Sociobiol. 64(4):529–539. 10.1007/s00265-009-0868-5

Rios-Soto J et al. 2017. Description of the distress call in *Espadarana prosoblepon* and the post-amplexus vocal display in *Centrolene savagei* (Anura: Centrolenidae). Herpetol Notes. 10:27–29.

Ritchie MG. 1992. Variation in male song and female preference within a population of *Ephippiger ephippiger* (Orthoptera: Tettigoniidae). Anim Behav. 43(5):845–855. 10.1016/S0003-3472(05)80207-6

Robertson JM, Lips KR, Heist EJ. 2008. Fine scale gene flow and individual movements among subpopulations of *Centrolene prosoblepon* (Anura: Centrolenidae). Rev Biol Trop. 56(1):13–26. 10.15517/rbt.v56i1.5506

Rodríguez-Correa V, González-Alzate S, Rada M, Vargas-Salinas F. 2024. The advertisement call of *Espadarana prosoblepon* (Anura: Centrolenidae) from a population in the Central Andes of Colombia. Phyllomedusa. 23(2):165–178. 10.11606/issn.2316-9079.v23i2p165-178

Rojas Montoya M, López-Aguirre Y, González-Acosta C, Vargas-Salinas F. 2024. Repertorio de señales acústicas en la rana de cristal *Hyalinobatrachium tatayoi* (Anura: Centrolenidae). Rev Latinoam Herpetol. 7(1):60–82. 10.22201/fc.25942158e.2024.1.747

RStudio Team. 2025. RStudio: Integrated Development Environment for R. http://www.posit.co/

Russo D, Jones G. 2002. Identification of twenty-two bat species (Mammalia: Chiroptera) from Italy by analysis of time-expanded recordings of echolocation calls. J Zool. 258(1):91–103. 10.1017/S0952836902001231

Ryan MJ. 1988. Constraints and patterns in the evolution of anuran acoustic communication. In: Fritzsch, B; Ryan, M J; Wilczynski, W; Hetherington, T E; Walkowiak, W, editor. The Evolution of The Amphibian Auditory System. Wiley; p 637–677.

Sakata JT, Hampton CM, Brainard MS. 2008. Social Modulation of Sequence and Syllable Variability in Adult Birdsong. J Neurophysiol. 99(4):1700–1711. 10.1152/jn.01296.2007

Searcy WA, Nowicki S. 2010. Reliability and Deception in Signaling Systems. In: The evolution of animal communication. Princeton University Press 10.1515/9781400835720

Teixeira BF da V, Zaracho VH, Giaretta AA. 2016. Advertisement and courtship calls of *Dendropsophus nanus* (Boulenger, 1889) (Anura: Hylidae) from its type locality (Resistencia, Argentina). Biota Neotrop. 16(4): e20160183. 10.1590/1676-0611-BN-2016-0183

Titus K, Mosher JA, Williams BK. 1984. Chance-corrected Classification for Use in Discriminant Analysis: Ecological Applications. Am Midl Nat. 111(1):1–7. 10.2307/2425535

Toledo LF et al. 2015. The anuran calling repertoire in the light of social context. Acta Ethol. 18(2):87–99. 10.1007/s10211-014-0194-4

Toledo LF, Haddad CFB. 2005. Acoustic Repertoire and Calling Behavior of *Scinax fuscomarginatus* (Anura, Hylidae). J Herpetol. 39(3):455–464. 10.1670/139-04A.1

Townsend SW, Rasmussen M, Clutton-Brock T, Manser MB. 2012. Flexible alarm calling in meerkats: the role of the social environment and predation urgency. Behav Ecol. 23(6):1360–1364. 10.1093/beheco/ars129

Vargas-Salinas F et al. 2014. Breeding and parental behaviour in the glass frog *Centrolene savagei* (Anura: Centrolenidae). J Nat Hist. 48(27–28):1689–1705. 10.1080/00222933.2013.840942

Welch AM, Semlitsch RD, Gerhardt HC. 1998. Call Duration as an Indicator of Genetic Quality in Male Gray Tree Frogs. Science. 280(5371):1928–1930. 10.1126/science.280.5371.1928

Wells KD. 1977. The Courtship of Frogs. In: Taylor, D.H., Guttman SI, editor. The Reproductive Biology of Amphibians. Springer US; p 233–262. 10.1007/978-1-4757-6781-0_7

Wells KD, Schwartz JJ. 2007. The Behavioral Ecology of Anuran Communication. In: Narins, P.M., Feng, A.S., Fay, R.R., Popper A., editor. Hearing and Sound Communication in Amphibians. 10.1007/978-0-387-47796-1_3

Wells KD, Taigen TL. 1986. The effect of social interactions on calling energetics in the gray treefrog (*Hyla versicolor*). Behav Ecol Sociobiol. 19(1):9–18. 10.1007/BF00303837

Zakon H, Oestreich J, Tallarovic S, Triefenbach F. 2002. EOD modulations of brown ghost electric fish: JARs, chirps, rises, and dips. J Physiol. 96(5–6):451–458. 10.1016/S0928-4257(03)00012-3

Zuberbühler K. 2008. Audience effects. Curr Biol. 18(5):R189–R190. 10.1016/j.cub.2007.12.041

